# Quantifying myelin density in the feline auditory cortex

**DOI:** 10.1101/2023.09.29.560251

**Authors:** Austin Robertson, Daniel J. Miller, Adam Hull, Blake E. Butler

**Affiliations:** Graduate program in Neuroscience, University of Western Ontario, London, ON, Canada; Department of Psychology, University of Western Ontario, London, ON, Canada; Undergraduate program in Neuroscience, University of Western Ontario, London, ON, Canada; Western Institute for Neuroscience, University of Western Ontario, London, ON, Canada; National Centre for Audiology, University of Western Ontario, London, ON, Canada

**Keywords:** Myelin, Auditory Cortex, Stereology, Feline

## Abstract

The cerebral cortex comprises many distinct regions that differ in structure, function, and patterns of connectivity. Current approaches to parcellating these regions often take advantage of functional neuroimaging approaches that can identify regions involved in a particular process with reasonable spatial resolution. However neuroanatomical biomarkers are also very useful in identifying distinct cortical regions either in addition to, or in place of functional measures. For example, differences in myelin density are thought to relate to functional differences between regions, are sensitive to individual patterns of experience, and have been shown to vary across functional hierarchies in a predictable manner. Accordingly, the current study provides quantitative stereological estimates of myelin density for each of the 13 regions that make up the feline auditory cortex. We demonstrate that significant differences can be observed between auditory cortical regions, with the highest myelin density observed in the regions that comprise the auditory core (i.e., the primary auditory cortex and anterior auditory field). Moreover, our myeloarchitectonic map suggests that myelin density varies in a hierarchical fashion that conforms to the traditional model of spatial organization in auditory cortex. Taken together, these results establish myelin as a useful biomarker for parcellating auditory cortical regions, and provide detailed estimates against which other, less invasive methods of quantifying cortical myelination may be compared.

## Introduction

Neuroscience has a longstanding interest in the structure-function relationships that define specific brain regions. Understanding the networks of areas that underlie complex behaviors is a common aim of cognitive neuroscientists, while identifying the underlying source of aberrant neural activity or behavior is critical to neurological assessment and therapeutic treatment. It is reasonable to propose that functionally distinct regions could be parcellated based on anatomical differences that reflect unique computational demands and patterns of connectivity. Accordingly, at the turn of the twentieth century at the direction of Oskar and Cecile Vogt, Brodmann published ‘Localization in the Cerebral Cortex’, in which he suggested that the brain could be divided into 52 areas based on differences in cytoarchitecture (Brodmann 1909). Impressively, this classic text provides many of the core principles that continue to guide attempts to resolve associations between cortical anatomy and function today (Annese 2009; Zilles 2018). For example, Brodmann’s delineations form the basis of at least one probabilistic map used in modern imaging analyses (e.g., the Talairach atlas), and his description of a universal 6-layer structure in the cerebral cortex is now universally accepted (Zilles 2018). Early *functional* maps of the cerebral cortex expanded upon these anatomical parcellations, often relying on lesion and/or neurosurgical studies from small numbers of patients. Recent technological advances including functional magnetic resonance imaging (fMRI), transcranial magnetic stimulation (TMS), and transcranial direct current stimulation (tDCS) have dramatically accelerated the development of more robust atlases. For example, fMRI has been used to dissociate regions that encode basic stimulus features (e.g., line fragments or particular orientations) from those that encode feature conjunctions and object representations in the visual cortical hierarchy (see Tootell et al. 1998 for review). Similarly, relatively brief fMRI sequences have been developed that allow one to delineate regions of the human auditory cortex that are organized by sound frequency from those that respond to more complex stimuli like vocalizations (e.g., Chevillet et al., 2011). These functional approaches have been instrumental in assigning functional roles to anatomically distinct regions of the brain.

fMRI-based parcellation typically involves collecting a pair of fMRI image series that, when contrasted, reveal the pattern of neural activity evoked by a stimulus feature of interest (e.g., the pattern of activity evoked by a stationary dot is subtracted from the activity evoked by that same dot when in motion to reveal motion-sensitive areas of the brain). While noninvasive imaging approaches are widespread, they suffer from a number of limitations. Firstly, it has been pointed out that the distribution of activity over the voxels within a region of interest may vary considerably depending on the context in which stimuli are presented, and the nature of the contrasts drawn (Friston et al. 2006; Poldrack 2007). As a result, parcellation via functional localizer scans may introduce bias and/or inappropriately constrain the definition of a particular region of interest (Friston et al., 2006). Moreover, functional localizer scans are of little use in identifying a region of interest when: 1) the response properties of the area are poorly defined or are highly similar to adjacent brain areas; or 2) the response properties of the region of interest are atypical (e.g., areas of deafferented sensory cortex). With these shortcomings in mind, there has been a renewed interest in examining markers that allow individual brain regions to be identified based on structural differences.

### Myeloarchitecture

There has been significant interest in myelin - a lipid-protein sheath that envelops axons of the central and peripheral nervous system - as a potential anatomical basis for cortical parcellation (e.g., Vogt & Vogt, 1919; Nieuwenhuys 2013). Within the central nervous system, myelin density is highest in white matter, with myelinated fibers extending into gray matter throughout the cortex (Vogt 1903). Generally, the density of myelinated fibers decreases from deep to superficial cortical layers; however, the absolute density of myelin across cortical depth can be highly heterogeneous across regions of the brain. These differences arise from local differences in cortical thickness, compactness of fiber layers, and the length and strength of radial fiber bundles (Nieuwenhuys 2013). Moreover, myelin density may differ across regions due to differences in the nature of the computations performed and/or patterns of connectivity to other areas of the brain. For example, myelin density has been shown to be inversely correlated with intracortical circuit complexity (Glasser et al. 2014). Accordingly, myelin gradients have been observed in typically developed sensory cortices, whereby core regions show higher levels of myelination than areas at subsequent steps along the functional hierarchies to which they project (Glasser and Van Essen 2011; Glasser et al. 2014; Miller et al. 2012; Miller et al. 2013). Since myelination may limit novel synapse formation and/or rearrangement, these hierarchical gradients may contribute to the observation that core sensory regions often show limited evidence of experience-dependent plasticity compared to higher-level regions that appear more responsive to ongoing sensory experience (Glasser et al. 2014; McGee et al. 2005). These myelination gradients are also in accordance with spatiotemporal patterns of development (Brody et al. 1987; Kinney et al. 1988) whereby sensory regions that develop earlier in life are myelinated earlier and more completely than regions that develop at a later age (Miller et al, 2012; Miller et al. 2013; Stadelmann et al. 2019).

Importantly, non-invasive methods have been developed that appear sensitive to the amount of myelin in different brain regions (see Lazari & Lipp 2021 for review). It is important to note, however, that these approaches typically provide only an indirect measure of myelination (see Lutti et al. 2014 for review). These methods typically involve MR imaging, and take advantage of the fact that myelin content affects relaxation rates (Mackay et al. 2009), magnetization transfer (Wolff & Balaban 1989), and the direction of local diffusion of water molecules within the myelin sheath (Beaulieu 2002). These approaches have historically been limited to estimating the myelin content of cubic mm-sized voxels within the brain; however recent advances in MRI sequences and multi-compartment modeling have improved the spatial resolution of non-invasive myelin quantification (Heath et al. 2018; Mezer et al. 2013). Thus, myelin density is expected to vary in a predictable manner across cortical regions, and these differences may be quantifiable both histologically and via non-invasive approaches across humans and animal models.

### Myelin as a biomarker of plasticity/disease

As a result of its role in accelerating action potential propagation, myelin has been shown to be integral in shaping and supporting large-scale functional neural networks (Hunt et al. 2016). Moreover, cortical myelination has been shown to be sensitive to early experience and dynamic across the lifespan, reflecting individual differences in experience, learning, and functional status (Flechsig 1901; Mount and Monje 2017). The onset of complex cognitive processes in humans occurs concurrently with the maturation of myelin-rich regions of the CNS (Williamson and Lyons 2018; Scantlebury et al. 2014; Mabbot et al. 2006). However, periods of social deprivation, particularly early in development, have been associated with a reduction of myelin content in the medial prefrontal cortex (Liu et al. 2012; Makinodan et al. 2012; Long et al. 2018). Additionally, significant changes in myelin density within the intraparietal sulcus have been observed following training on a complex, visually-guided motor skill (McKenzie et al., 2014; Scholz et al. 2009), while impairing the development and differentiation of oligodendrocytes significantly impairs learning (Xiao et al., 2016; McKenzie et al., 2014) and retention (Kaller et al. 2017).

Atypical myelination is observed in a number of conditions associated with disrupted cortical function including Multiple Sclerosis (Rahmanzadeh et al. 2021), schizophrenia (Takahashi et al. 2011), Amyotrophic Lateral Sclerosis, and Huntington’s disease (Huang et al. 2015). Moreover, a growing body of literature suggests Alzheimer’s disease is associated with atypical patterns of myelination (Ota et al. 2019), and that changes in myeloarchitecture may correlate with the onset of Alzheimer’s symptomatology (Maitre et al. 2023). The clear relationships between myeloarchitecture and experience-dependent plasticity/disease etiology (Fields 2005; 2008) underscore the important role that myelin quantification may play in tracking the anatomical status of cortical regions, surgical planning, and in predicting associated functional change following therapeutic interventions.

### The present study: Quantifying myelination of auditory cortex

In order to assess the potential of myelin density to parcellate cortical subregions within an established functional hierarchy, the current study sought to quantify patterns of myelination across the auditory cortex of the cat. The cat provides a simplified, tractable model of the primate auditory cortex (Lomber and Malhotra, 2008) within which a functional hierarchy has been proposed (Lee & Winer, 2011). Due in part to the size and accessibility of the feline auditory cortex (∼20% of the cortical volume, much of which is exposed on the cortical surface), the cat has emerged as an important model of auditory cortical structure and function to which early foundational work in visual cortical organization can be compared. While there has been some historical disagreement on the parcellation of the feline auditory cortex, we adhere to a popular model which suggests it comprises 13 distinct regions (Winer & Schreiner, 2011). Additionally, different names have been ascribed to these regions over time; in order to maintain continuity between the present study and our previous studies describing patterns of connectivity to auditory cortex (e.g., Chabot et al. 2015; Butler et al. 2016; 2017; 2018), we describe two core areas (A1, AAF), three additional low-level regions that are tonotopically organized (PAF, VPAF, VAF), five higher-order regions that respond to more complex acoustic features (A2, fAES, DZ, IN, T) and three multisensory regions that integrate auditory and non-auditory perceptual cues (dPE, iPE, vPE). These regions can be further divided into two parallel streams of processing: a dorsal stream specialized for the representation of auditory space, and a ventral stream specialized in spectral analysis. This type of functional division within the auditory cortex was first proposed based on electrophysiological recordings in nonhuman primates (e.g., Rauschecker & Tian 2000; Tian et al. 2001), and has since been extended to the feline auditory cortex (Figure 1), where detailed functional hierarchies have been proposed based on patterns of thalamocortical projections to, and corticocortical projections between regions in each stream (Lee & Winer 2011; see Winer & Schreiner 2011 for a summary of the evidence for functional streams). These ‘connectional fingerprints’ have been effectively used to delineate cortical regions and infer functional hierarchies across a number of species (e.g., macaque: Stephan et al. 2001; human: Behrens & Johansen-Berg 2005), and have been shown to correlate with changes in both cytoarchitecture and function in the feline auditory cortex (Rose & Woolsey 1948). Here, we aim to 1) quantify regional differences in myelin density in feline auditory cortex; and 2) examine the extent to which myelin density is related to the position of a given region within the functional hierarchy of sound processing.

**Figure 1.**
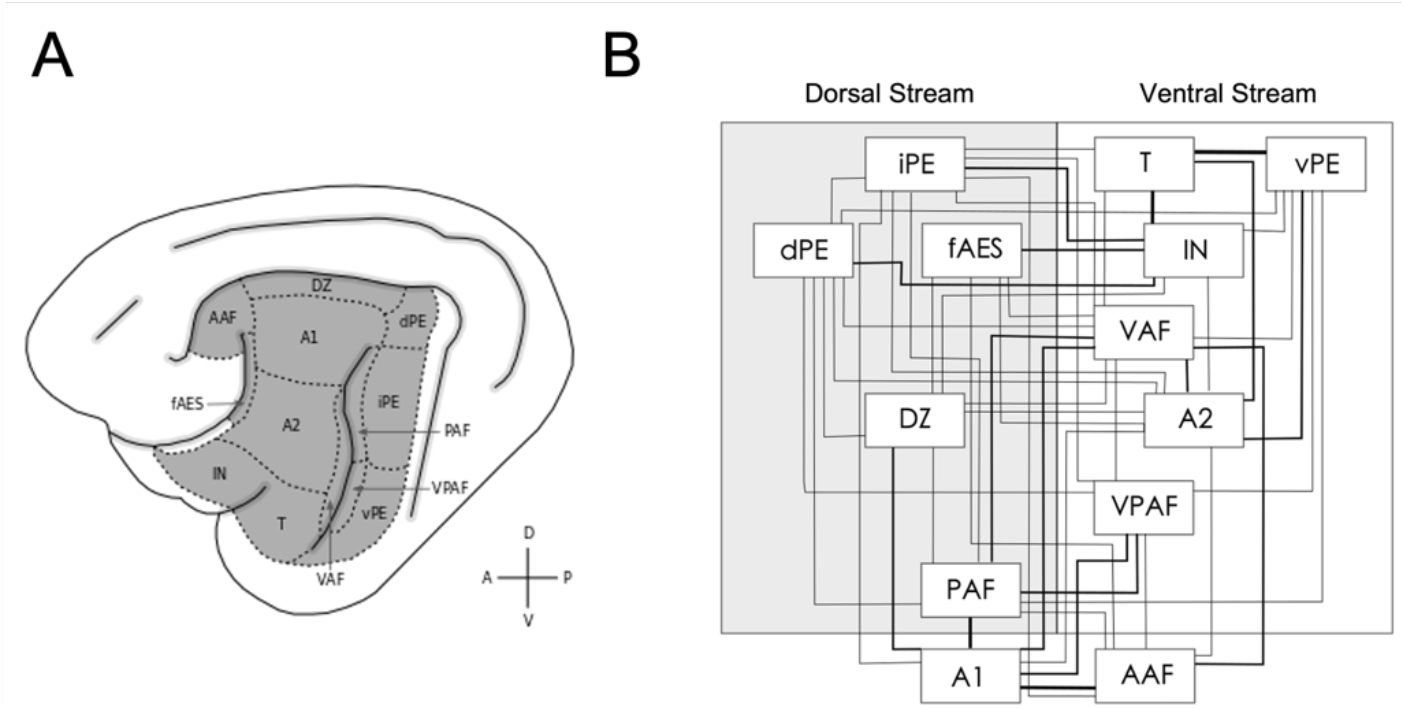
A. Lateral view of the feline brain highlighting auditory cortical regions. B. Hierarchical arrangement of these regions (adapted from Lee & Winer 2011). A1: primary auditory cortex; AAF: anterior auditory field; PAF: posterior auditory field; VPAF: ventral posterior auditory field; DZ: dorsal zone; A2: second auditory cortex; VAF: ventral auditory field; dPE/iPE/vPE: dorsal/intermediate/ventral divisions of posterior ectosylvian gyrus; fAES: auditory division of the anterior ectosylvian sulcus; IN: insular area; T: temporal area.

## Methods

Patterns of myelinated fiber length density (MFLD) were examined in three male domestic short-haired cats obtained from a USDA licensed commercial breeding facility (Marshall Bioresources; Waverly, NY). At the time of sacrifice, each animal was anesthetized with ketamine (4 mg/kg, i.m.) and dexdomitor (0.03 mg/kg, i.m.) and a catheter was inserted into the cephalic vein. Sodium pentobarbital (40 mg/kg, i.v.) was administered to induce general anesthesia, and heparin (1 mL: anticoagulant) and 1% sodium nitrite (1 mL: vasodilator) were administered intravenously. Each animal was then transcardially perfused with physiological saline (1 L), followed by 4% paraformaldehyde (2 L), and 10% sucrose (2L). Following perfusion, brains were blocked in the coronal plane between Horsley-Clarke levels A18 and P2 (a region spanning the entirety of auditory cortex; Horsley & Clarke, 1908), removed, and immersed in a 30% sucrose solution until sunk for cryoprotection (∼1 week). Brains were then sectioned in the coronal plane at a thickness of 60 um using a cryostat (Thermo-Fisher, Waltham, Mass). Six series were collected at 360 um intervals: two series were stained for myelin using a modified Gallyas Silver Impregnation method (Gallyas 1979, Pistorio et al. 2006, Joseph et al. 2019); one series was processed using the monoclonal antibody SMI-32 (Biolegend; Sternberger & Sternberger 1983) to allow for laminar and areal delineation of auditory cortical subfields; one series was stained for Nissl bodies using cresyl violet to assist with delineation; and two series served as spaced to be processed as necessary.

### SMI-32 Immunohistochemistry

Antibodies sensitive to SMI-32 recognize a non-phosphorylated epitope of neurofilament proteins, a ligand that is differentially expressed across regions of the auditory cortex, allowing them to be delineated (Mellot et al. 2010). To visualize patterns of SMI-32 reactivity, sections were rinsed with 0.1M physiological buffer (PB; 3 x 5 min), and endogenous peroxidase was blocked with 0.5% hydrogen peroxide in 70% ethanol (30 min). Sections were then rinsed with 0.1 M PB (4 x 5 min), incubated in 5% normal goat serum in 0.1M PB (45 min), and incubated in mouse-anti-SMI-32 antibody (Vector Laboratories, Burlingame, CA, USA; 1/2000 in 2% NGS in 0.1M PB) overnight. The following day, sections were rinsed with 0.1 M PB (4 x 5 min) and incubated in biotinylated goat anti-mouse IgG antibody (Vector Laboratories: 1/200 in 2% NGS in 0.1M PB) for 30 min. Sections were then rinsed with 0.1 M PB (3 x 10 min), and incubated in an avidin–biotin–horseradish peroxidase solution (Vectastain Elite ABC, Vector Laboratories) for 90 minutes. Sections were then rinsed with 0.1 M PB (3 x 10 mins), and incubated with a diaminobenzene–nickel chromogen solution for 15 min. Finally, sections were rinsed with 0.1 M PB (3 x 5 min), mounted from 0.01 M PB on gelatinized slides, air-dried, cleared, and coverslipped.

### Myelin Staining

The silver impregnation method (Gallyas 1979; Hackett 2001; Miller et al. 2012, 2013) allows for the visualization of individual myelinated fibers in cortical tissue. While other methods are available to visualize myelin, the resolution provided by the Gallyas silver stain is much greater than that of the myelin basic protein antibody staining method, the Luxol Fast Blue method, Weil’s rapid method, or Weigert’s stain (Pistorio et al. 2006). Specifically, the modified Gallyas protocol used in the present study has been shown to be sensitive to small diameter myelinated fibers that are difficult to visualize with alternative methods. To stain for myelin, sections were rinsed in deionized water (3 x 5 min), and then incubated in a 2:1 solution of pyridine and acetic anhydride (30 minutes). Sections were then rinsed in 0.5% acetic acid (3 x 3 min), and incubated in an Ammoniacal Silver solution (0.1% ammonium nitrate, 0.1% silver nitrate w/v; 60 min). Sections were subsequently rinsed in 0.5% acetic acid (3 x 3 min) and incubated in a developer solution containing equal parts sodium carbonate anhydrous (5% w/v), and a solution of ammonium nitrate (0.2% w/v), silver nitrate (0.2% w/v), and tungstosilicic acid hydrate (1% w/v) for 5 minutes. Sections were then rinsed with deionized water (3 x 5 min) and bleached in potassium ferricyanide solution (0.5% w/v; 3 min) before being rinsed again in deionized water (3 x 5 min). This process of development, bleaching, and rinsing was repeated twice to ensure the tissue was well-developed and uniformly stained. At that point, tissue was fixed by incubating in sodium thiosulfate (0.5% w/v; 5 min), and rinsed thoroughly in deionized water (3×10 min). Sections were mounted from gel alcohol (gelatine [1% w/v] in equal parts deionized water and 95% alcohol) on gelatinized slides, air dried, cleared, and coverslipped with permount prior to counting.

### Myelin Quantification

Mounted sections were examined using a Zeiss AxioImager M2 microscope equipped with a Lumina HR camera. First, regions of interest (ROIs) throughout auditory cortex were defined for each animal based on cytoarchitecture and sulcal/gyral landmarks defining the borders between adjacent areas. SMI-32 has a high affinity for a dephosphorylated epitope on the medium- and high-molecular weight subunits of neurofilament proteins, which results in robust labeling of pyramidal cells and dendrites. The specific patterns of labelling across the cortical mantle differ noticeably between adjacent regions of auditory cortex, (Mellott et al. 2010) such that borders can be readily identified on SMI-32-labelled tissue. Additionally, the patterns of labelling observed in auditory cortical areas differ from those observed in visual and somatosensory cortices (van der Gucht et al., 2001) such that the outer limits of auditory cortex can also be readily identified. Using these patterns of SMI-32 expression, each of the 13 regions of auditory cortex were identified and outlined using the contour feature in StereoInvestigator (MBF Bioscience; Williston, VT). Each region was then further divided into supragranular (layers I-III), granular (layer IV), and infragranular layers (layers V-VI) for a total of 39 ROIs in each animal. These ROIs were then transferred to adjacent sections stained for myelin to define the boundaries within which myelin estimates were drawn. Myelinated fiber length density (MFLD) was computed as the length of myelinated fiber found within a given volume of tissue, and estimates for each ROI were generated using the Spaceballs probe (Mouton et al. 2002) in StereoInvestigator (MBF Bioscience; Williston, VT) under Koehler illumination at 100x. The Spaceballs probe has been used extensively to generate unbiased length estimates based on the probability that the surface of a spherical or hemispherical volume of tissue is intersected by a linear feature (e.g., myelinated axon). This approach represents a significant advance over previous sampling techniques; for example, estimates of fiber number obtained using a 2-dimensional cross-section of tissue are biased because the number of profiles observed is unrelated to the number of fibers in the tissue by any known function, and no correction factor can resolve the associated sampling bias (Gunderson et al., 1988). Sampling parameters for the current study are provided in Table 1, and were selected to ensure that between 70-100 probes would be evaluated within each ROI across a minimum of 3 sections (Schmitz & Hof 2005). Hemispheric Spaceballs probes were placed in a systematic random fashion across each ROI, and the points at which myelinated fibers intersected the surface of the probe were marked. To convert myelin length estimates generated by Stereoinvestigator to length density estimates, the total myelin length obtained for each ROI was divided by the total tissue volume of that ROI. Tissue volumes were estimated by Stereoinvestigator using the Cavalieri principle (Gundersen & Jensen, 1987) by multiplying the total area of the ROI by the mean mounted thickness which was measured at every 5th sampling location. Finally, the coefficient of error for each estimate was computed to ensure that the variability of the estimate was sufficiently low to ensure an accurate stereological estimate (Schmitz & Hof 2005; see Appendix A).

**Table 1.** Average number of pyramidal somata per 500 um in auditory cortex (from Mellott et al. 2010).

|  | A1 | PAF | DZ | fAES | AAF | VPAF | A2 | VAF | IN | T |
| --- | --- | --- | --- | --- | --- | --- | --- | --- | --- | --- |
| Layer II | 26.3 | 18.2 | 34.7 | 27.3 | 38.2 | 0.0 | 28.7 | 27.5 | 0.0 | 0.0 |
| Layer III | 40.7 | 32.6 | 58.7 | 36.8 | 54.4 | 0.0 | 38.6 | 33.6 | 0.0 | 0.0 |
| Layer V | 38.1 | 35.7 | 48.2 | 32.8 | 54.8 | 19.7 | 49.5 | 30.2 | 29.2 | 22.8 |
| Layer VI | 38.6 | 15.6 | 38.6 | 15.9 | 34.5 | 0.0 | 31.5 | 23.5 | 11.4 | 0.0 |

\*Regions along the posterior ectosylvian gyrus (dPE/iPE/vPE) were not examined in this study
\*\*No somata were counted in layers I/IV

### Statistical Analysis

To allow for comparison across regions of interest of different sizes, myelin length estimates computed for each ROI were divided by the total volume of that ROI to generate myelin length density measures. Myelin length density values were then normalized to an animal’s mean MFLD across regions to account for variance that may arise due to differences in the developing/bleaching procedures across animals (although, the modified protocol used in the current study has been shown to reduced this variability considerably compared to the original Gallyas protocol [Pistorio et al. 2006]). Normalized myelin length density values were then compared using a repeated-measures ANOVA with region (13 levels; Figure 1) and depth (infragranular, granular, supragranular) treated as within-subjects factors. Tukey’s honestly significant difference test was used to examine differences in myelin density across cortical depths while correcting for multiple comparisons (familywise a = 0.05).

To examine how myelin length density varied between regions across the entire thickness of cortex, volume-weighted estimates for each region were computed by multiplying the MFLD observed at each depth by the proportional volume of the ROI at that depth. For example, the myelin density of A1 was computed as follows:

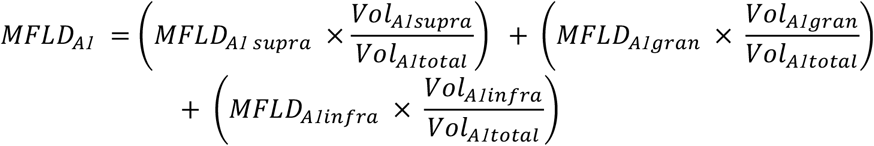

| Abbreviation | Definition |
| --- | --- |
| MFLD<sub>A1</sub> | Myelinated fiber length density (MFLD) of primary auditory cortex (A1) |
| MFLD<sub>A1supra</sub> | MFLD in supragranular A1 |
| MFLD<sub>A1gran</sub> | MFLD in granular A1 |
| MFLD<sub>A1infra</sub> | MFLD in infragranular A1 |
| Vol<sub>A1supra</sub> | Volume of supragranular A1 |
| Vol<sub>A1gran</sub> | Volume of granular A1 |
| Vol<sub>A1infra</sub> | Volume of infragranular A1 |

Volume-weighted myelin length density estimates were compared across regions using a repeated-measures ANOVA with region (13 levels) treated as a within-subjects factor. Tukey’s honestly significant difference test was used to examine pairwise differences in myelin density while correcting for multiple comparisons.

## Results

The current study sought to quantify and compare the patterns of myelination across the feline auditory cortex (Figure 2). Regions of interest were created that comprised the infragranular, granular, and supragranular layers of each of the 13 regions of auditory cortex (for a total of 39 ROIs), and stereological estimates of myelin length density were computed for each ROI. Figure 3 shows the MFLD of each region and cortical depth for each animal examined, as well as the mean MFLD across animals. For ease of visualization, the 13 regions have been arranged according to their position within the proposed functional hierarchy of processing in auditory cortex (Lee & Winer 2011), with dorsal stream regions on the left and ventral stream regions on the right. As expected, significant differences in myelin density were observed across cortical depth (F(2,4)=111.83, p<0.01); infragranular layers were more densely myelinated that the granular layer (q=0.354, p<0.001), which was more densely myelinated than the supragranular layers (q=0.616, p<0.001).

**Figure 2.**
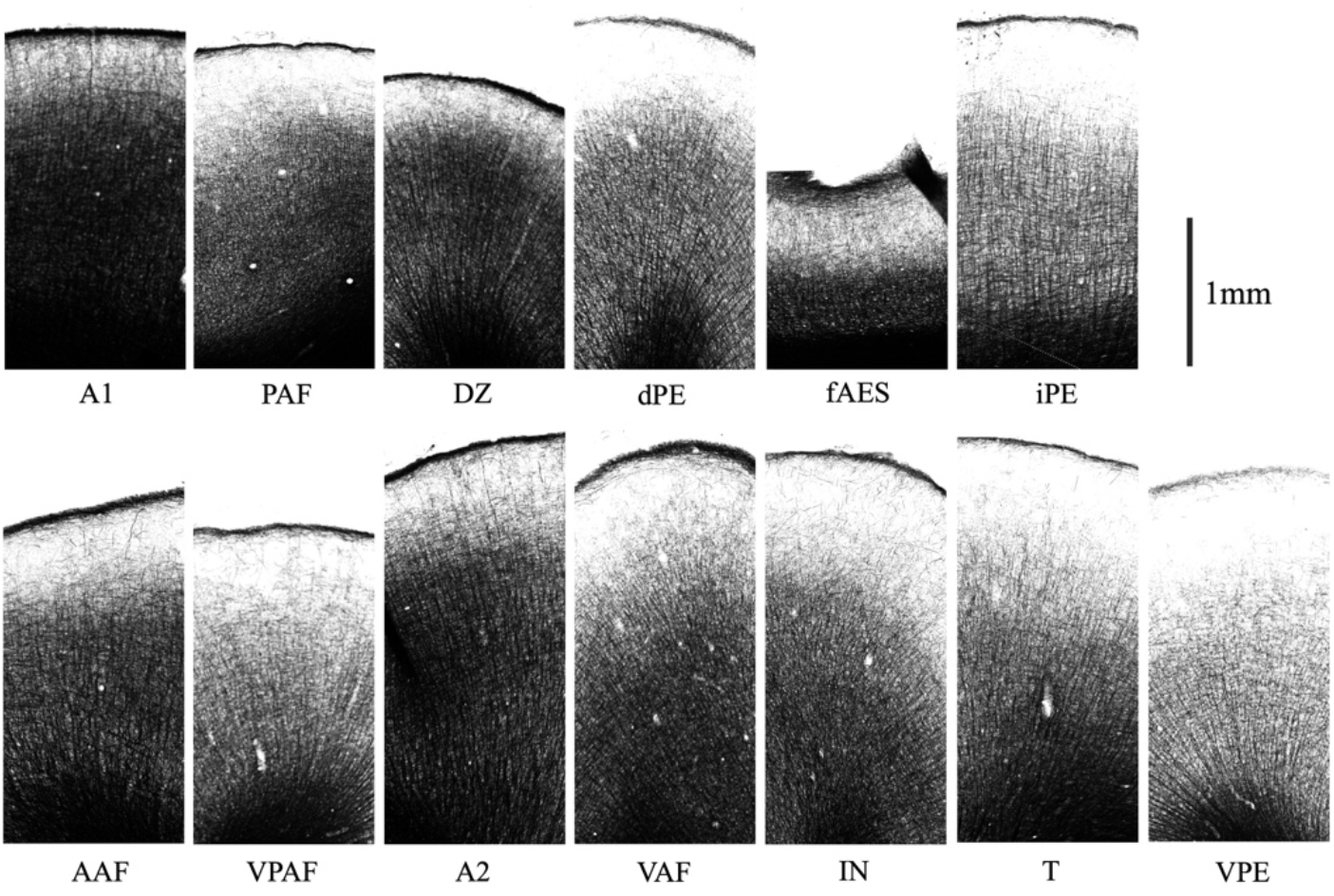
Photomicrographs of cortical myelination across the regions of the feline auditory cortex. Regions are arranged by position within the putative functional hierarchy of processing in auditory cortex (Lee & Winer 2011), with dorsal stream regions on the top, and ventral stream regions on the bottom. Within each image, the cortical surface is positioned at top, with the white matter boundary at bottom. Scale bar shows 1mm.

**Figure 3.**
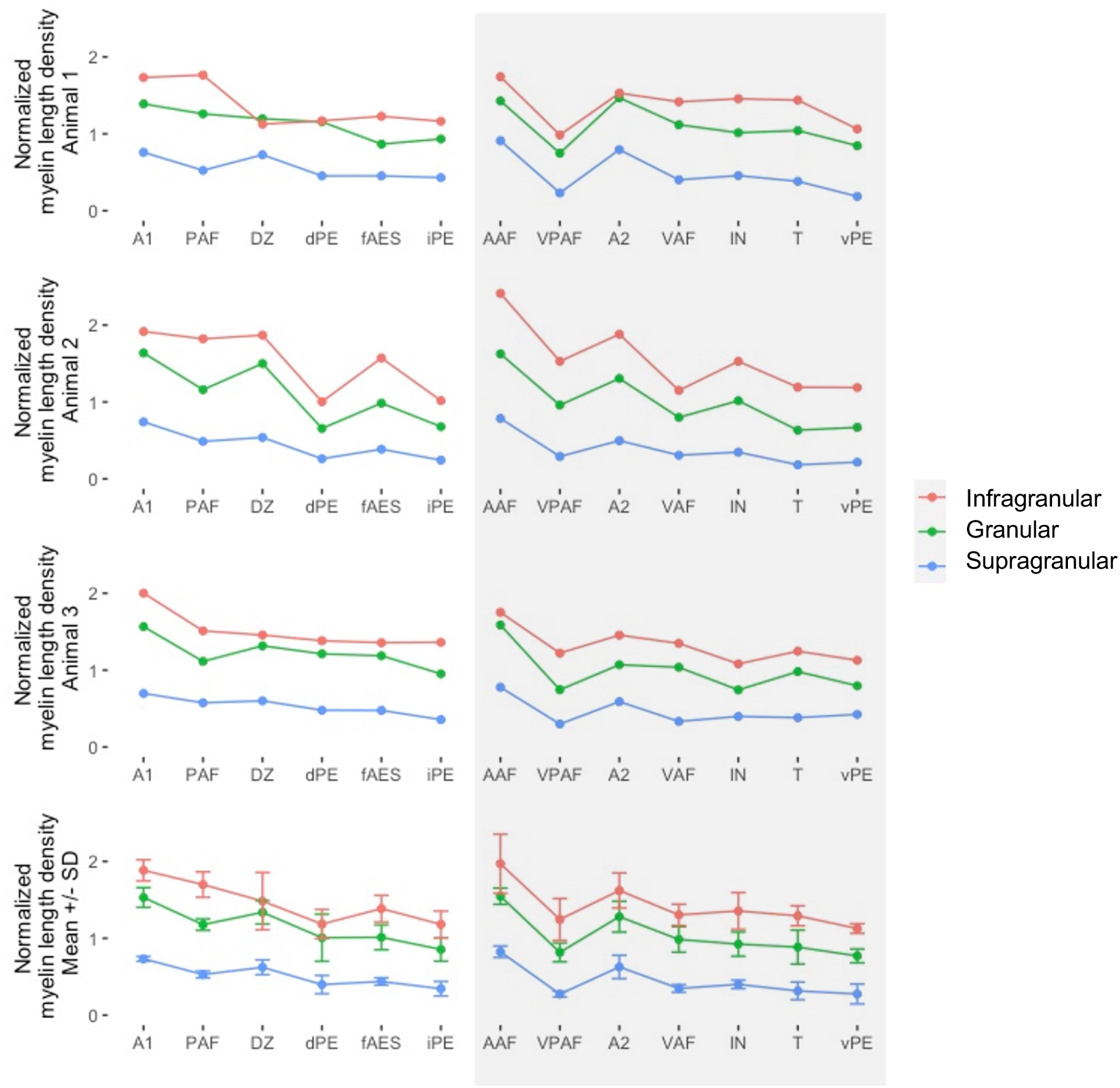
Myelin length density for each auditory cortical region of interest, at each depth examined (red = infragranular, green = granular, blue = supragranular). Regions are arranged by position within the putative functional hierarchy of processing in auditory cortex (Lee & Winer 2011), with dorsal stream regions on the left, and ventral stream regions on the right. Individual animal values (top three panels) were normalized to the mean density across regions within-animal to account for variance that may arise due to differences in the developing/bleaching procedures across animals. A significant depth effect was observed, with myelin density increasing with cortical depth across all regions of interest examined. Error bars in bottom panel show the standard error of the mean.

In the current study, the effect of depth did not depend on which region of auditory cortex was considered (F(24,48)=1.18, p=0.31). Accordingly, data were collapsed across depths to examine regional differences in myelin density. To do so, a single volume-weighted estimate of myelin density was computed for each region (Figure 4). A repeated-measures ANOVA using these volume-weighted estimates revealed a significant effect of region (F(12,24)=12.53, p<0.001). A complete list of pairwise comparisons between regions is provided in Appendix B. In general, myelin density decreased from core to higher level regions within each stream of the putative functional hierarchy. Indeed, differences in myelin density most frequently reached statistical significance when core regions (primary auditory cortex [A1], anterior auditory field [AAF]) were compared to high level areas (e.g., dorsal, intermediate, and ventral divisions of the posterior ectosylvian gyrus [dPE, iPE, vPE], insular and temporal auditory areas [IN,T], the auditory division of the anterior ectosylvian sulcus [fAES]). Interestingly, despite being placed near the base of the ventral stream functional hierarchy, the ventral posterior auditory field (VPAF) appeared to be less densely myelinated than higher-level areas - an observation that was consistent across animals.

**Figure 4.**
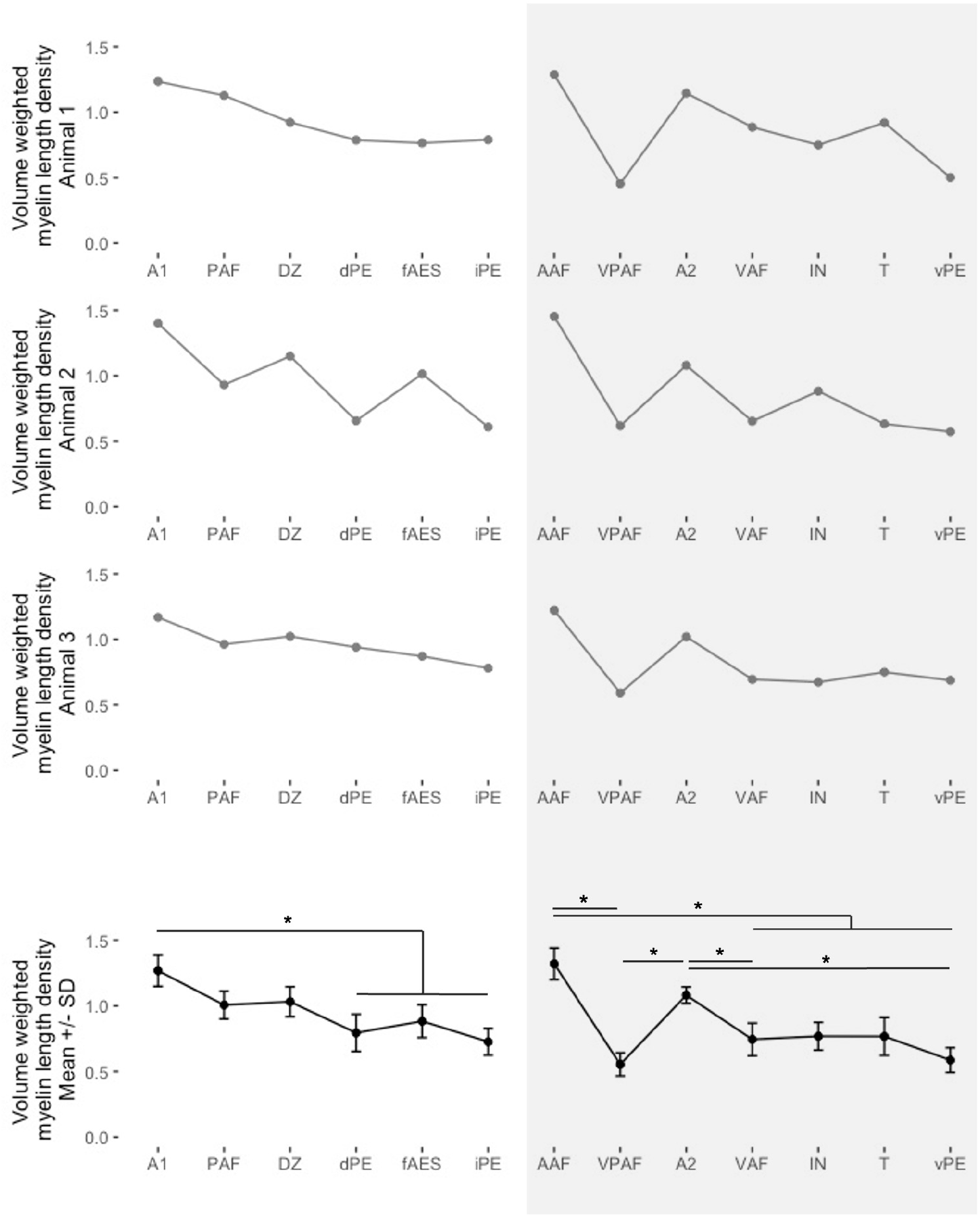
Stereological estimates of myelinated fiber length density (MFLD) for each auditory cortical region of interest. Regions are arranged by position within the putative functional hierarchy of processing in auditory cortex (Lee & Winer 2011), with dorsal stream regions on the left, and ventral stream regions on the right. Error bars in bottom panel show the standard error of the mean. Significant differences in myelin length density within the same stream are shown (* denotes p < 0.05, corrected for multiple comparisons [familywise *α* = 0.05]); see appendix A for a summary of all pairwise comparisons.

## Discussion

This study presents the first detailed quantification and comparison of myelin density across the feline auditory cortex. As expected, MFLD increased with cortical depth across all regions; the deepest layers of cortex, which are most proximal to the white matter, are more densely myelinated than more superficial layers. In human cortex, differences in the relative myelin density between layers occasionally create sharp borders that distinguish adjacent regions (e.g., between divisions of the frontal lobe [Vogt & Vogt 1919]). However, a vast majority of human cortical borders are marked by transitions in myelination that are subtler and more gradual (Von Economo & Koskinas 1925). This in accordance with the gradual shifts in cytoarchitecture that are often observed between cortical areas (e.g., most borders between auditory cortical regions were described as ‘transitional’ by Mellott and colleagues [2010]). In the current study, borders between adjacent brain regions were often apparent based on overall myelin density (Figure 2); however, differences in relative myelin density across depths were rather uniform across auditory cortex. This was somewhat unexpected given previous accounts of laminar differences in the distribution of myelin, including between core and non-core regions of the primate auditory cortex (e.g., Hackett et al., 2001; see Niewenhuys 2013 for a comprehensive review of human myeloarchitectural variability). It is possible that some subtle variability in the laminar profile of myelination across the feline auditory cortex might emerge from an analysis approach that contrasts myelin density in individual layers or subdivisions of layers – the level granularity that distinguishes the types of myeloarchitectonic layering proposed by Vogt (1910) and others.

### Regional differences in myelin density

Auditory cortex has been traditionally defined as the portion of the brain that receives its primary thalamic inputs from the dorsal, medial, or ventral divisions of the medial geniculate nucleus (Hackett 2008). Within auditory cortex, distinct regions can be identified based on their patterns of thalamocortical and corticocortical connectivity, differences in cytoarchitecture, and unique functional properties. Based on these differences, the feline auditory cortex can be segmented into 13 regions (Winer & Schreiner 2011). In the current study, myelin density varied across these regions in a manner that was highly consistent across animals. Even at low magnification, many of the borders between adjacent regions could be identified based on shifts in myelin density.

The regional differences in myelin density observed here conform to the organizational structure that has been observed across mammalian auditory cortex: one or more ‘core’ regions adjoined by three secondary areas that comprise a ‘belt’ (AL, ML, CL), which further project to two tertiary regions comprising a ‘parabelt’ (rostral and caudal PB; Hackett 2008; Miller et al. 2013). While the core-belt-parabelt structure has been most commonly associated with detailed studies of primate auditory cortex (e.g., Kaas and Hackett 2000), a similar pattern of organization has been proposed based on patterns of corticocortical connectivity observed in the feline brain (Lee and Winer 2008a). It has been previously observed that the regions surrounding the auditory core including A2, DZ, fAES, and PAF show similar, belt-like patterns of thalamocortical (Lee and Winer 2008b) and corticocortical connectivity (Lee and Winer 2008a). Moreover, it was noted that areas along the rostral portion of the posterior ectosylvian gyrus (dPE, iPE, vPE) and at the ventral extent of auditory cortex (In, T) show parabelt-like patterns of projections, with little to no input from core areas (Lee and Winer, 2008a). The current study is in accordance with these connectional data, and suggests that myelin density decreases as one moves from putative core to belt, to parabelt regions of auditory cortex (Figure 5). While similar patterns have been observed previously, the current work extends upon these findings by quantifying differences between cortical areas have largely been described qualitatively (but, see Miller et al., 2012; 2013). In doing so, we are able to quantify the consistency across animals, both across the cortical mantle and across regions of auditory cortex. The existence of a heavily myelinated auditory core is also in accordance with neuroimaging studies of the human auditory cortex (Sigalovsky et al. 2006; Dick et al., 2012; Lutti et al. 2013). Thus, the patterns observed in the current study provide additional support for the feline auditory cortex as a tractable model of the structure and function of human auditory cortex. Moreover, patterns of myelin density may aid in identifying homologous regions across species (Miller et al. 2013), and in aligning cortical maps (e.g., Glasser & Van Essen, 2011).

**Figure 5.**
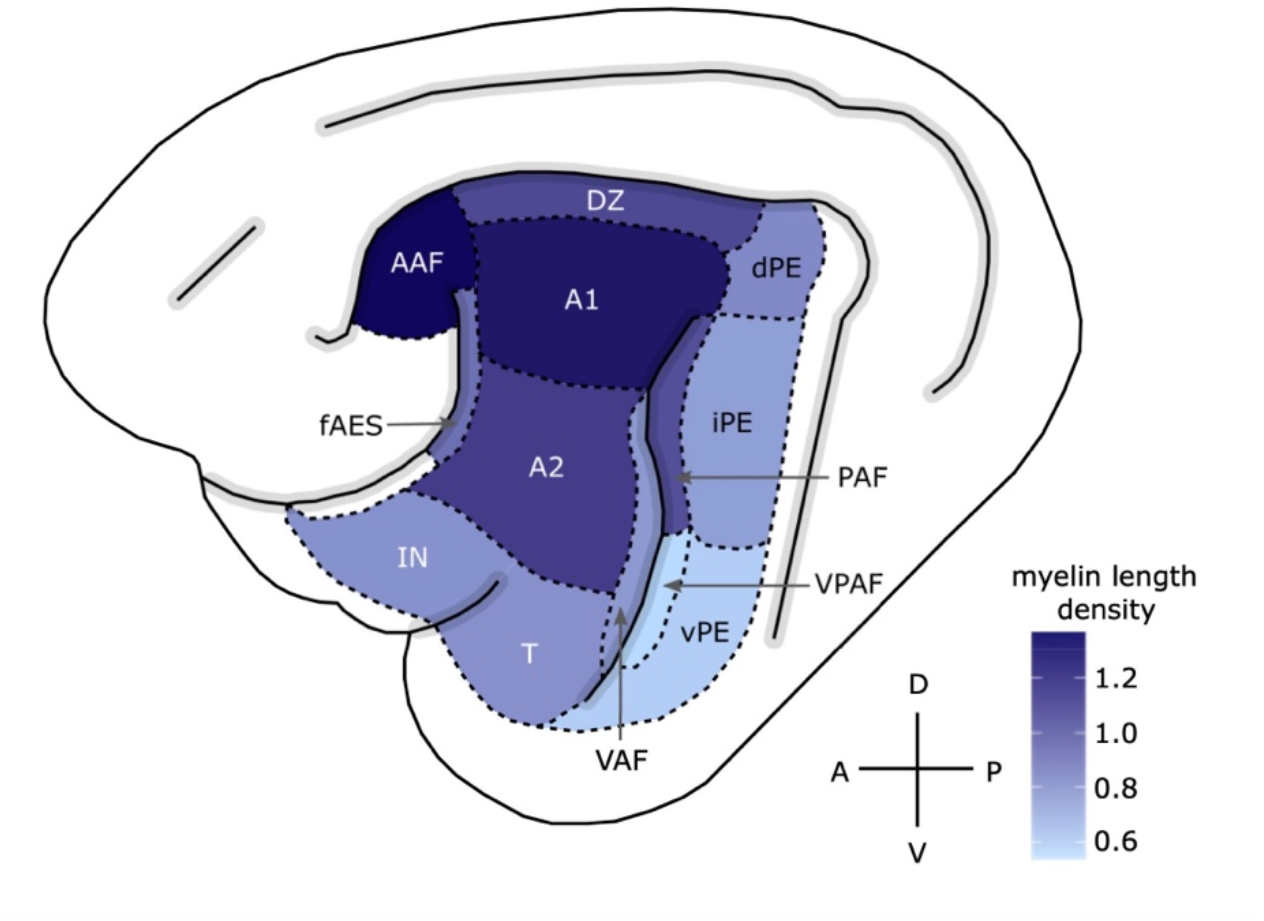
Lateral view of the cat brain showing how myelin length density changes across auditory cortex. Darker colors show regions of denser MFLD (see inset colorbar). AAF: anterior auditory field; fAES: auditory division of the anterior ectosylvian sulcus; DZ: dorsal zone of auditory cortex; A1: primary auditory cortex; A2: second auditory cortex; IN: insular area; T: temporal area; PAF; posterior auditory field; VAF: ventral auditory field; VPAF: ventral posterior auditory field; dPE: dorsal division of the posterior ectosylvian gyrus; iPE: intermediate area of the posterior ectosylvian gyrus; vPE: ventral division of the posterior ectosylvian gyrus.

### Hierarchical gradient

The core-belt-parabelt organization of auditory cortex establishes a functional hierarchy wherein ascending thalamocortical projections project predominantly to core regions, which in turn project to higher-level (belt, parabelt) areas (Rauschecker et al. 1997). Based on patterns of connectivity, the feline auditory cortex has been further arranged into two parallel functional hierarchies that process information related to sound location (dorsal stream) and spectral composition (ventral stream; Lee & Winer 2011; Figure 1). Based on previous work documenting gradients in myelin density within sensory cortices (e.g., Miller et al. 2013; Dick et al. 2017), the current study predicted that myelin density would decrease with increasing hierarchical position along each of these pathways. Indeed, with the exception of one region (the ventral posterior auditory field; VPAF), the pattern of myelin density observed in the current study conformed to the proposed arrangement.

In part, the hierarchical gradient of myelin density presented here likely reflects regional differences in neuronal density (and thus, differences in the number of myelinated axons per unit volume; see Table 1). However, regional differences may also be related to the time course of neural development, and may play an important role in regional differences in experience-dependent plasticity (Fields 2005; 2008). As a general principle, myelination ascends along a hierarchical order of increasing functional complexity (Yakovlev & Lecours 1967), and axons in regions that are myelinated early are also myelinated more rapidly and more completely (Stadelmann et al. 2019). Since core regions of auditory cortex are the principal targets of thalamocortical afferents that form the earliest cortical circuitry (Yakovlev & Lecours 1967; Taymourtash et al. 2023), it stands to reason they would be most densely myelinated.

The early development and dense myelination of core auditory cortical areas relative to higher-level regions may contribute to differences in neuroplastic potential across the functional hierarchy. For example, there is evidence that the dorsal zone (DZ; Lomber et al. 2010), posterior auditory field (PAF; Lomber et al. 2010), and auditory field of the anterior ectosylvian sulcus (fAES; Meredith et al. 2011) take on visual functions following profound, early-onset hearing loss; however, evidence for functional reorganization of core feline auditory regions remains contested (Stewart and Starr 1970; Rebillard et al. 1977; Kral et al. 2003). The lack of robust evidence for reorganization in core auditory areas may be related to their myelin density, which has been suggested to play a role in stabilizing existing intracortical circuits by inhibiting the growth of axons and formation of new synapses (Chen et al. 2000; Kapfhammer and Schwab 1994; Glasser et al. 2014). Accordingly, higher-level areas that are more sparsely myelinated may also retain a greater ability to adapt to novel experiences.

### Myelin as a marker for non-invasive segmentation

Regional differences in myelin density follow a predictable hierarchy, and many of the borders between regions are readily identifiable based on their relative myelination. Accordingly, the current study supports the idea that myelin density may be an effective biomarker for segmenting functionally distinct auditory cortical areas. Fortunately, non-invasive means of quantifying myelin have received tremendous attention over the past decade. For example, magnetic resonance imaging-based approaches that purport to be sensitive to myelin include T1w/T2w imaging, myelin water fraction imaging, and various indices based on magnetization transfer imaging (e.g., magnetization transfer ratio, magnetization transfer saturation) and diffusion imaging, among others (see Lazari and Lipp 2021 for review). MRI-based approaches to quantifying myelin rely upon proxy measures (e.g., differential rates of relaxation, inferences drawn from the spin properties or motion of water molecules). While many of these measures have been shown to correlate with histological estimates, methodological variance across studies and evidence of publication bias have obscured direct comparisons across approaches (Lazari and Lipp 2021). By quantifying myelin density across auditory cortex, the current study provides detailed histological estimates to which different, less invasive methods can be compared and validated.

## Conclusion

Advances in noninvasive quantification of neuroanatomical features has renewed interests in generating detailed histological maps. Myeloarchitectonic maps are of particular interest, as differences in myelin density have been observed within and between cortical regions, and are associated with functional status. Moreover, changes in myelination have been associated with developmental trajectories, patterns of experience, and a number of neurological disorders. The current study provides detailed qualifications of myelin density across the feline auditory cortex - 13 functionally distinct regions that form two parallel hierarchies of processing. Regional differences in myelination are in line with the classical core-belt-parabelt organization of auditory cortex, and reveal a hierarchical pattern in which lower-level regions are more densely myelinated than higher-order areas. The results presented here provide important insights into the anatomical organization of auditory cortex, and confirm that quantifiable differences in myelin density between brain regions are reliably observed. Moreover, these detailed histological quantifications provide ground-truth estimates against which other, less invasive measures of myelin density may be compared. Overall, the current study highlights the potential of myelin density as a biomarker for cortical parcellation, and for the study of how the anatomy and physiology of specific brain regions are affected by individual experience.

## Acknowledgements

The authors extend thanks to Katelyn Kittel for animal care support, and to Dr. Faraj Haddad for support with protocol development.

## Declarations

### Funding

This work received financial support from the Natural Sciences and Engineering Research Council and from BrainsCAN - an initiative funded by the Canada First Research Excellence Fund.

### Competing Interests

The authors have no relevant financial or non-financial interests to disclose.

## Author Contributions

**Austin Robertson:** Formal Analysis, Investigation, Writing - Original Draft, Visualization. **Daniel J. Miller:** Conceptualization, Methodology, Investigation, Writing - Review & Editing, Supervision. **Adam Hull:** Investigation. **Blake E. Butler:** Conceptualization, Writing - Review & Editing, Supervision, Project Administration, Funding Acquisition.

## Data Availability

Data generated during the current study are available on the Open Science Framework (https://osf.io/24jb6/).

## Ethics Approval

All surgical and experimental procedures were conducted in accordance with the Canadian Council on Animal Care’s *Guide to the Care and Use of Experimental Animals* (Olfert et al. 1993) and were approved by the University of Western Ontario Animal Care Committee.

## Appendix A

Coefficients of error in myelin length density estimates for each ROI in each animal.

| Region | Depth | Animal 1 CoE | Animal 2 CoE | Animal 3 CoE |
| --- | --- | --- | --- | --- |
| A1 | Infragranular | 0.04 | 0.04 | 0.05 |
|  | Granular | 0.05 | 0.05 | 0.06 |
|  | Supragranular | 0.04 | 0.04 | 0.05 |
| PAF | Infragranular | 0.04 | 0.05 | 0.05 |
|  | Granular | 0.05 | 0.06 | 0.05 |
|  | Supragranular | 0.05 | 0.05 | 0.05 |
| DZ | Infragranular | 0.05 | 0.04 | 0.06 |
|  | Granular | 0.06 | 0.06 | 0.06 |
|  | Supragranular | 0.05 | 0.05 | 0.06 |
| dPE | Infragranular | 0.06 | 0.04 | 0.05 |
|  | Granular | 0.06 | 0.05 | 0.06 |
|  | Supragranular | 0.07 | 0.06 | 0.05 |
| fAES | Infragranular | 0.05 | 0.06 | 0.05 |
|  | Granular | 0.07 | 0.06 | 0.05 |
|  | Supragranular | 0.05 | 0.05 | 0.06 |
| iPE | Infragranular | 0.05 | 0.04 | 0.04 |
|  | Granular | 0.06 | 0.06 | 0.05 |
|  | Supragranular | 0.06 | 0.06 | 0.05 |
| AAF | Infragranular | 0.05 | 0.06 | 0.05 |
|  | Granular | 0.06 | 0.06 | 0.06 |
|  | Supragranular | 0.07 | 0.06 | 0.05 |
| VPAF | Infragranular | 0.05 | 0.06 | 0.08 |
|  | Granular | 0.07 | 0.07 | 0.06 |
|  | Supragranular | 0.09 | 0.07 | 0.07 |
| A2 | Infragranular | 0.05 | 0.07 | 0.05 |
|  | Granular | 0.05 | 0.06 | 0.05 |
|  | Supragranular | 0.05 | 0.06 | 0.06 |
| VAF | Infragranular | 0.09 | 0.08 | 0.07 |
|  | Granular | 0.08 | 0.08 | 0.08 |
|  | Supragranular | 0.09 | 0.08 | 0.06 |
| IN | Infragranular | 0.04 | 0.05 | 0.05 |
|  | Granular | 0.05 | 0.06 | 0.07 |
|  | Supragranular | 0.06 | 0.06 | 0.06 |
| T | Infragranular | 0.04 | 0.06 | 0.05 |
|  | Granular | 0.05 | 0.08 | 0.05 |
|  | Supragranular | 0.06 | 0.1 | 0.06 |
| vPE | Infragranular | 0.04 | 0.05 | 0.06 |
|  | Granular | 0.06 | 0.06 | 0.06 |
|  | Supragranular | 0.07 | 0.07 | 0.06 |

## Appendix B

Full output of Tukey’s HSD test showing all pairwise comparisons of weighted myelin length density estimates. * p<0.05; ** p<0.01; ***p<0.001

**Appendix B.** Full output of Tukey's HSD test showing all pairwise comparisons of weighted myelin length density estimates. \* p<0.05; \*\* p<0.01; \*\*\*p<0.001
| Region 1 | Region 2 | q | p |
| --- | --- | --- | --- |
| A1 | PAF | -0.261 | 0.25 |
|  | DZ | -0.236 | 0.383 |
|  | dPE | -0.473 | 0.0014 ** |
|  | fAES | -0.384 | 0.0149 * |
|  | iPE | -0.541 | 0.000218 *** |
|  | AAF | 0.0525 | 1 |
|  | VPAF | -0.713 | 0.0000023 *** |
|  | A2 | -0.186 | 0.715 |
|  | VAF | -0.523 | 0.000359 *** |
|  | IN | -0.499 | 0.00069 *** |
|  | T | -0.5 | 0.000671 *** |
|  | vPE | -0.681 | 0.00000534 *** |
| PAF | DZ | 0.0249 | 1 |
|  | dPE | -0.212 | 0.538 |
|  | fAES | -0.123 | 0.975 |
|  | iPE | -0.28 | 0.171 |
|  | AAF | 0.313 | 0.0842 |
|  | VPAF | -0.452 | 0.00244 ** |
|  | A2 | 0.0747 | 1 |
|  | VAF | -0.262 | 0.245 |
|  | IN | -0.238 | 0.37 |
|  | T | -0.239 | 0.364 |
|  | vPE | -0.42 | 0.00589 ** |
| DZ | dPE | -0.237 | 0.376 |
|  | fAES | -0.148 | 0.912 |
|  | iPE | -0.305 | 0.101 |
|  | AAF | 0.289 | 0.145 |
|  | VPAF | -0.477 | 0.00124 ** |
|  | A2 | 0.0498 | 1 |
|  | VAF | -0.287 | 0.15 |
|  | IN | -0.263 | 0.24 |
|  | T | -0.264 | 0.236 |
|  | vPE | -0.445 | 0.00302 ** |
| dPE | fAES | 0.0888 | 0.998 |
|  | iPE | -0.0681 | 1 |
|  | AAF | 0.526 | 0.000333 *** |
|  | VPAF | -0.24 | 0.358 |
|  | A2 | 0.287 | 0.15 |
|  | VAF | -0.0498 | 1 |
|  | IN | -0.0259 | 1 |
|  | T | -0.0269 | 1 |
|  | vPE | -0.207 | 0.571 |
| fAES | iPE | -0.157 | 0.878 |
|  | AAF | 0.437 | 0.00372 ** |
|  | VPAF | -0.329 | 0.0587 |
|  | A2 | 0.198 | 0.634 |
|  | VAF | -0.139 | 0.943 |
|  | IN | -0.115 | 0.986 |
|  | T | -0.116 | 0.985 |
|  | vPE | -0.296 | 0.123 |
| iPE | AAF | 0.594 | 0.0000526 *** |
|  | VPAF | -0.172 | 0.801 |
|  | A2 | 0.355 | 0.0312 * |
|  | VAF | 0.0183 | 1 |
|  | IN | 0.0423 | 1 |
|  | T | 0.0412 | 1 |
|  | vPE | -0.139 | 0.941 |
| AAF | VPAF | -0.766 | 0.000000616 *** |
|  | A2 | -0.239 | 0.367 |
|  | VAF | -0.576 | 0.0000862 *** |
|  | IN | -0.552 | 0.000165 *** |
|  | T | -0.553 | 0.00016 *** |
|  | vPE | -0.733 | 0.0000014 *** |
| VPAF | A2 | 0.527 | 0.000319 *** |
|  | VAF | 0.19 | 0.687 |
|  | IN | 0.214 | 0.523 |
|  | T | 0.213 | 0.53 |
|  | vPE | 0.0328 | 1 |
| A2 | VAF | -0.337 | 0.0488 * |
|  | IN | -0.313 | 0.0852 |
|  | T | -0.314 | 0.0833 |
|  | vPE | -0.494 | 0.000782 *** |
| VAF | IN | 0.0239 | 1 |
|  | T | 0.0229 | 1 |
|  | vPE | -0.158 | 0.875 |
| IN | T | -0.00101 | 1 |
|  | vPE | -0.182 | 0.745 |
| T | vPE | -0.181 | 0.751 |

## Notes

https://osf.io/24jb6/

### Competing Interest Statement

The authors have declared no competing interest.

### Summary of Updates

This version has been revised following peer review. Some additional references to existing work in primates has been added, and a justification for pursuing this work in a feline model has been added.

